## Supplementary Materials for "Why we publish where we do: Faculty publishing values and their relationship to review, promotion and tenure expectations"

**Meredith T. Niles^1*^, Lesley A. Schimanski^2^, Erin C. McKiernan^3^, Juan P. Alperin^4*^**

1 Department of Nutrition and Food Sciences & Food Systems Program, University of Vermont, Burlington VT 05405

2 Scholarly Communications Lab, Simon Fraser University, Vancouver, Canada

3 Departamento de Física, Universidad Nacional Autónoma de México, Mexico City, Mexico

4 Scholarly Communications Lab, Simon Fraser University, Vancouver, Canada

^*^Corresponding authors

| Supplementary Table 1. Mean responses, standard error (SE) and p values for publishing decisions and productivity by tenure status. ANOVA were used for statistical significance tests. | | | | | |
| --- | --- | --- | --- | --- | --- |
| **Variable** | **Tenured** | **SE** | **Non-tenure** | **SE** | **p=** |
| pubs published | 3.15 | 0.92 | 3.24 | 0.91 | 0.467 |
| merit pay | 1.91 | 1.43 | 2.04 | 1.55 | 0.597 |
| readership | 5.03 | 1.27 | 5.02 | 1.23 | 0.989 |
| Journal IF | 4.18 | 1.41 | 4.61 | 1.58 | 0.029 |
| society journal | 3.47 | 1.66 | 3.45 | 1.73 | 0.958 |
| journal read | 4.49 | 1.30 | 4.50 | 1.41 | 0.927 |
| journal peers read | 4.66 | 1.36 | 4.76 | 1.44 | 0.599 |
| journal citations | 3.80 | 1.49 | 4.11 | 1.45 | 0.126 |
| journal prestige | 4.70 | 1.17 | 4.95 | 1.23 | 0.123 |
| open access | 3.23 | 1.62 | 3.50 | 1.51 | 0.216 |
| journal cost | 3.70 | 1.80 | 3.71 | 1.82 | 0.976 |

| Supplementary Table 2. Mean responses, standard error (SE) and p values for publishing decisions and productivity by gender and institution type. ANOVA were used for statistical significance tests. | | | | | | | | | | |
| --- | --- | --- | --- | --- | --- | --- | --- | --- | --- | --- |
| **Variable** | **Female** | **SE** | **Male** | **SE** | **p=** | **R Type** | **SE** | **M Type** | **SE** | **p=** |
| pubs published | 3.05 | 0.88 | 3.28 | 0.96 | 0.022 | 3.30 | 0.89 | 2.89 | 0.93 | 0.000 |
| merit pay | 2.08 | 1.66 | 1.87 | 1.29 | 0.308 | 2.03 | 1.50 | 1.77 | 1.36 | 0.256 |
| readership | 5.00 | 1.25 | 5.02 | 1.30 | 0.927 | 5.11 | 1.16 | 4.82 | 1.47 | 0.073 |
| Journal IF | 4.33 | 1.45 | 4.26 | 1.49 | 0.721 | 4.38 | 1.42 | 4.13 | 1.57 | 0.199 |
| society journal | 3.54 | 1.77 | 3.41 | 1.58 | 0.543 | 3.51 | 1.61 | 3.39 | 1.82 | 0.596 |
| journal read | 4.59 | 1.31 | 4.38 | 1.38 | 0.197 | 4.57 | 1.26 | 4.29 | 1.50 | 0.109 |
| journal peers read | 4.69 | 1.45 | 4.66 | 1.34 | 0.853 | 4.73 | 1.37 | 4.54 | 1.43 | 0.308 |
| journal citations | 3.94 | 1.48 | 3.84 | 1.48 | 0.595 | 3.94 | 1.51 | 3.74 | 1.45 | 0.310 |
| journal prestige | 4.73 | 1.25 | 4.76 | 1.15 | 0.812 | 4.79 | 1.24 | 4.65 | 1.10 | 0.363 |
| open access | 3.24 | 1.59 | 3.38 | 1.60 | 0.488 | 3.24 | 1.55 | 3.43 | 1.69 | 0.386 |
| journal cost | 4.14 | 1.86 | 3.36 | 3.36 | 0.001 | 3.80 | 1.78 | 3.51 | 1.83 | 0.232 |

| Supplementary Table 3. Spearman's correlations and p values for publishing decisions and productivity by age | | |
| --- | --- | --- |
| **Variable** | **Age** | **p =** |
| pubs published | -0.109 | 0.064 |
| merit pay | 0.080 | 0.256 |
| readership | -0.026 | 0.662 |
| Journal IF | -0.156 | 0.009 |
| society journal | 0.124 | 0.039 |
| journal read | -0.024 | 0.690 |
| journal peers read | -0.089 | 0.140 |
| journal citations | -0.182 | 0.002 |
| journal prestige | -0.165 | 0.005 |
| open access | 0.097 | 0.109 |
| journal cost | -0.091 | 0.149 |

| Supplementary Table 4. Spearman's correlations and p values for perception of the RPT process by age | | |
| --- | --- | --- |
| **Variable** | **Age** | **p value** |
| rpt blog | 0.160 | 0.010 |
| rpt book chapter | 0.122 | 0.045 |
| rpt book | -0.011 | 0.852 |
| rpt pub numbers | -0.026 | 0.664 |
| rpt performance | 0.317 | 0.000 |
| rpt media | -0.046 | 0.459 |
| rpt pre print | -0.037 | 0.581 |
| rpt open access | 0.250 | 0.000 |
| rpt society | 0.104 | 0.092 |
| rpt journal IF | -0.080 | 0.189 |
| rpt journal name | -0.065 | 0.281 |
| rpt pub total | -0.058 | 0.332 |

| Supplementary Table 5. Mean responses and p values for perception of the RPT process by tenure status. ANOVA were used for statistical significance tests. | | | | | |
| --- | --- | --- | --- | --- | --- |
|  | **Tenured** | **SE** | **Non-tenure** | **SE** | **p=** |
| rpt blog | 1.86 | 1.10 | 1.74 | 0.95 | 0.227 |
| rpt book chapter | 3.61 | 1.39 | 3.14 | 1.26 | 0.009 |
| rpt book | 4.34 | 1.56 | 3.74 | 1.58 | 0.025 |
| rpt pub numbers | 5.32 | 0.94 | 5.22 | 1.15 | 0.391 |
| rpt performance | 2.20 | 1.57 | 2.19 | 1.57 | 0.978 |
| rpt media | 3.07 | 1.51 | 3.03 | 1.50 | 0.597 |
| rpt pre print | 2.30 | 1.35 | 2.18 | 1.34 | 0.552 |
| rpt open access | 2.21 | 1.40 | 1.88 | 1.21 | 0.072 |
| rpt society | 3.56 | 1.55 | 3.21 | 1.53 | 0.075 |
| rpt journal IF | 4.59 | 1.37 | 4.85 | 1.37 | 0.521 |
| rpt journal name | 3.84 | 1.21 | 4.82 | 1.33 | 0.600 |
| rpt pub total | 3.36 | 0.99 | 5.51 | 0.88 | 0.419 |

| Supplementary Table 6. Mean responses and p values for perception of the RPT process by gender and institution type. ANOVA were used for statistical significance tests. | | | | | | | | | | |
| --- | --- | --- | --- | --- | --- | --- | --- | --- | --- | --- |
| **Variable** | **Female** | **SE** | **Male** | **SE** | **p=** | **R Type** | **SE** | **M Type** | **SE** | **p=** |
| rpt blog | 1.89 | 1.12 | 1.76 | 1.00 | 0.322 | 1.79 | 1.03 | 1.90 | 1.14 | 0.480 |
| rpt book chapter | 3.63 | 1.36 | 3.35 | 1.38 | 0.106 | 3.29 | 1.30 | 3.86 | 1.42 | 0.001 |
| rpt book | 4.35 | 1.49 | 3.98 | 1.66 | 0.060 | 4.07 | 1.58 | 4.38 | 1.59 | 0.139 |
| rpt pub numbers | 5.51 | 0.88 | 5.11 | 1.05 | 0.001 | 5.35 | 0.98 | 5.19 | 1.03 | 0.220 |
| rpt performance | 2.22 | 1.50 | 2.19 | 1.63 | 0.931 | 2.17 | 1.51 | 2.24 | 1.65 | 0.773 |
| rpt media | 3.11 | 1.53 | 3.04 | 1.49 | 0.746 | 3.02 | 1.45 | 3.04 | 1.56 | 0.939 |
| rpt pre print | 2.29 | 1.38 | 2.26 | 1.33 | 0.852 | 2.14 | 1.28 | 2.64 | 1.47 | 0.043 |
| rpt open access | 2.06 | 1.39 | 2.17 | 1.33 | 0.525 | 2.01 | 1.19 | 2.30 | 1.58 | 0.102 |
| rpt society | 3.58 | 1.66 | 3.34 | 1.44 | 0.214 | 3.49 | 1.46 | 3.46 | 1.73 | 0.892 |
| rpt journal IF | 4.82 | 1.38 | 4.50 | 1.35 | 0.057 | 4.81 | 1.25 | 4.37 | 1.54 | 0.014 |
| rpt journal name | 4.92 | 1.24 | 4.75 | 1.25 | 0.265 | 4.97 | 1.12 | 4.58 | 1.39 | 0.013 |
| rpt pub total | 5.60 | 0.82 | 5.20 | 1.05 | 0.001 | 5.45 | 0.88 | 5.30 | 1.11 | 0.212 |

| Supplementary Table 7. Ordered logistic model predicting merit pay as a factor in publishing decisions (Model 1). Total n= 154. | | | | | | |
| --- | --- | --- | --- | --- | --- | --- |
| **Variable** | **Odds Ratio** | **Std. Err** | **z** | **P value** | **95% confidence interval** | |
| age | 1.268 | 0.221 | 1.36 | 0.174 | 0.901 | 1.785 |
| gender | 0.812 | 0.290 | -0.58 | 0.559 | 0.403 | 1.635 |
| r-type | 1.671 | 0.659 | 1.30 | 0.193 | 0.771 | 3.620 |
| tenured | 0.590 | 0.259 | -1.20 | 0.230 | 0.249 | 1.396 |
| pubs published | 1.067 | 0.202 | 0.34 | 0.730 | 0.737 | 1.545 |
| rpt pub numbers | 0.657 | 0.154 | -1.79 | 0.073 | 0.414 | 1.040 |
| rpt preprint | 1.083 | 0.156 | 0.55 | 0.581 | 0.816 | 1.436 |
| rpt open access | 1.355 | 0.190 | 2.16 | 0.031 | 1.028 | 1.784 |
| rpt society | 1.045 | 0.124 | 0.37 | 0.711 | 0.828 | 1.319 |
| rpt journal IF | 1.039 | 0.163 | 0.24 | 0.807 | 0.764 | 1.414 |
| rpt journal name | 0.928 | 0.157 | -0.44 | 0.659 | 0.666 | 1.293 |
| rpt pub total | 1.305 | 0.322 | 1.08 | 0.281 | 0.805 | 2.117 |

| Supplementary Table 8. Ordered logistic model predicting readership respondents want to reach as a factor in publishing decisions (Model 2). Total n= 203. | | | | | | | | | | | | |
| --- | --- | --- | --- | --- | --- | --- | --- | --- | --- | --- | --- | --- |
| **Variable** | | **Odds Ratio** | **Std Err** | | | **z** | | **P value** | | **95% confidence interval** | | |
| age | | 0.934 | 0.139 | | | -0.46 | | 0.648 | | 0.697 | | 1.252 |
| gender | | 1.135 | 0.339 | | | 0.43 | | 0.671 | | 0.632 | | 2.039 |
| r-type | | 0.905 | 0.292 | | | -0.31 | | 0.757 | | 0.480 | | 1.705 |
| tenured | | 1.273 | 0.460 | | | 0.67 | | 0.503 | | 0.628 | | 2.583 |
| pubs published | | 1.679 | 0.291 | | | 2.99 | | 0.003 | | 1.196 | | 2.359 |
| rpt pub numbers | | 1.104 | 0.189 | | | 0.58 | | 0.561 | | 0.790 | | 1.544 |
| rpt preprint | | 1.096 | 0.127 | | | 0.79 | | 0.430 | | 0.873 | | 1.375 |
| rpt open access | | 1.365 | 0.168 | | | 2.54 | | 0.011 | | 1.073 | | 1.737 |
| rpt society | | 0.956 | 0.088 | | | -0.49 | | 0.623 | | 0.798 | | 1.144 |
| rpt journal IF | | 0.806 | 0.104 | | | -1.68 | | 0.094 | | 0.627 | | 1.037 |
| rpt journal name | | 1.252 | 0.176 | | | 1.60 | | 0.110 | | 0.950 | | 1.650 |
| rpt pub total | | 1.318 | 0.232 | | | 1.57 | | 0.117 | | 0.933 | | 1.861 |
| Supplementary Table 9. Ordered logistic model predicting Journal Impact Factor as a factor in publishing decisions (Model 3). Total n= 203. | | | | | | | | | | | | |
| **Variable** | **Odds Ratio** | | | **Std Err** | **z** | | **P value** | | **95% confidence interval** | | | |
| age | 0.911 | | | 0.124 | -0.69 | | 0.492 | | 0.697 | | 1.190 | |
| gender | 0.905 | | | 0.250 | -0.36 | | 0.717 | | 0.527 | | 1.555 | |
| r-type | 0.853 | | | 0.256 | -0.53 | | 0.596 | | 0.474 | | 1.536 | |
| tenured | 0.605 | | | 0.214 | -1.42 | | 0.155 | | 0.302 | | 1.209 | |
| pubs published | 1.320 | | | 0.204 | 1.80 | | 0.072 | | 0.976 | | 1.787 | |
| rpt pub numbers | 1.066 | | | 0.172 | 0.39 | | 0.693 | | 0.777 | | 1.462 | |
| rpt preprint | 1.246 | | | 0.137 | 2.00 | | 0.045 | | 1.005 | | 1.545 | |
| rpt open access | 0.986 | | | 0.109 | -0.12 | | 0.902 | | 0.794 | | 1.226 | |
| rpt society | 0.969 | | | 0.084 | -0.36 | | 0.716 | | 0.818 | | 1.148 | |
| rpt journal IF | 1.884 | | | 0.249 | 4.79 | | 0.000 | | 1.454 | | 2.442 | |
| rpt journal name | 0.939 | | | 0.132 | -0.45 | | 0.653 | | 0.713 | | 1.236 | |
| rpt pub total | 0.907 | | | 0.153 | -0.58 | | 0.565 | | 0.652 | | 1.263 | |

| Supplementary Table 10. Ordered logistic model predicting society journals as a factor in publishing decisions (Model 4). Total n= 204. | | | | | | |
| --- | --- | --- | --- | --- | --- | --- |
| **Variable** | **Odds Ratio** | **Std Err** | **z** | **P value** | **95% confidence interval** | |
| age | 1.373 | 0.186 | 2.34 | 0.019 | 1.053 | 1.789 |
| gender | 1.026 | 0.277 | 0.10 | 0.924 | 0.604 | 1.743 |
| r-type | 1.277 | 0.379 | 0.82 | 0.410 | 0.714 | 2.285 |
| tenured | 0.698 | 0.235 | -1.07 | 0.286 | 0.361 | 1.351 |
| pubs published | 1.127 | 0.166 | 0.81 | 0.419 | 0.844 | 1.504 |
| rpt pub numbers | 0.795 | 0.138 | -1.32 | 0.187 | 0.566 | 1.118 |
| rpt preprint | 0.995 | 0.101 | -0.05 | 0.962 | 0.816 | 1.213 |
| rpt open access | 1.065 | 0.115 | 0.58 | 0.561 | 0.861 | 1.317 |
| rpt society | 1.720 | 0.160 | 5.83 | 0.000 | 1.433 | 2.064 |
| rpt journal IF | 0.914 | 0.113 | -0.73 | 0.465 | 0.718 | 1.164 |
| rpt journal name | 1.101 | 0.153 | 0.69 | 0.491 | 0.838 | 1.446 |
| rpt pub total | 1.167 | 0.192 | 0.94 | 0.348 | 0.845 | 1.613 |

| Supplementary Table 11. Ordered logistic model predicting journal/venue/publisher they regularly read as a factor in publishing decisions (Model 5). Total n= 205. | | | | | | | | | | | |
| --- | --- | --- | --- | --- | --- | --- | --- | --- | --- | --- | --- |
| **Variable** | | **Odds Ratio** | | **Std Err** | | **z** | **P value** | **95% confidence interval** | | | |
| age | | 1.057 | | 0.147 | | 0.40 | 0.688 | 0.806 | | 1.388 | |
| gender | | 0.825 | | 0.227 | | -0.70 | 0.486 | 0.481 | | 1.417 | |
| r-type | | 1.207 | | 0.363 | | 0.63 | 0.531 | 0.670 | | 2.175 | |
| tenured | | 0.767 | | 0.258 | | -0.79 | 0.430 | 0.397 | | 1.482 | |
| pubs published | | 1.248 | | 0.182 | | 1.52 | 0.128 | 0.938 | | 1.661 | |
| rpt pub numbers | | 1.009 | | 0.163 | | 0.05 | 0.957 | 0.734 | | 1.386 | |
| rpt preprint | | 1.072 | | 0.114 | | 0.65 | 0.515 | 0.870 | | 1.320 | |
| rpt open access | | 1.063 | | 0.116 | | 0.56 | 0.576 | 0.858 | | 1.317 | |
| rpt society | | 1.052 | | 0.091 | | 0.59 | 0.554 | 0.889 | | 1.246 | |
| rpt journal IF | | 0.843 | | 0.097 | | -1.49 | 0.135 | 0.673 | | 1.055 | |
| rpt journal name | | 1.309 | | 0.165 | | 2.15 | 0.032 | 1.024 | | 1.675 | |
| rpt pub total | | 0.980 | | 0.155 | | -0.13 | 0.900 | 0.719 | | 1.337 | |
| Supplementary Table 12. Ordered logistic model predicting journal/venue/publisher their peers regularly read as a factor in publishing decisions (Model 6). Total n= 203. | | | | | | | | | | | |
| **Variable** | **Odds Ratio** | | **Std Err** | | **z** | | **P value** | | **95% confidence interval** | | |
| age | 0.876 | | 0.122 | | -0.95 | | 0.341 | | 0.667 | | 1.151 |
| gender | 1.314 | | 0.368 | | 0.98 | | 0.329 | | 0.759 | | 2.276 |
| r-type | 0.971 | | 0.288 | | -0.10 | | 0.922 | | 0.543 | | 1.738 |
| tenured | 0.753 | | 0.262 | | -0.82 | | 0.414 | | 0.381 | | 1.488 |
| pubs published | 0.982 | | 0.148 | | -0.12 | | 0.901 | | 0.731 | | 1.319 |
| rpt pub numbers | 1.001 | | 0.162 | | 0.01 | | 0.994 | | 0.729 | | 1.374 |
| rpt preprint | 0.884 | | 0.094 | | -1.15 | | 0.248 | | 0.718 | | 1.090 |
| rpt open access | 1.114 | | 0.123 | | 0.98 | | 0.326 | | 0.898 | | 1.382 |
| rpt society | 1.055 | | 0.092 | | 0.61 | | 0.543 | | 0.888 | | 1.253 |
| rpt journal IF | 0.891 | | 0.107 | | -0.96 | | 0.336 | | 0.703 | | 1.128 |
| rpt journal name | 1.376 | | 0.185 | | 2.38 | | 0.018 | | 1.058 | | 1.791 |
| rpt pub total | 1.146 | | 0.183 | | 0.85 | | 0.393 | | 0.838 | | 1.568 |

| Supplementary Table 13. Ordered logistic model predicting how often a journal is cited as a factor in publishing decisions (Model 7). Total n= 203. | | | | | | |
| --- | --- | --- | --- | --- | --- | --- |
| **Variable** | **Odds Ratio** | **Std Err** | **z** | **P value** | **95% confidence interval** | |
| age | 0.829 | 0.113 | -1.37 | 0.170 | 0.634 | 1.084 |
| gender | 1.069 | 0.285 | 0.25 | 0.804 | 0.634 | 1.802 |
| r-type | 0.747 | 0.219 | -0.99 | 0.320 | 0.421 | 1.327 |
| tenured | 1.148 | 0.399 | 0.40 | 0.691 | 0.581 | 2.269 |
| pubs published | 1.107 | 0.161 | 0.70 | 0.485 | 0.832 | 1.473 |
| rpt pub numbers | 0.794 | 0.135 | -1.36 | 0.175 | 0.569 | 1.108 |
| rpt preprint | 1.258 | 0.134 | 2.15 | 0.031 | 1.021 | 1.551 |
| rpt open access | 0.930 | 0.099 | -0.68 | 0.494 | 0.755 | 1.145 |
| rpt society | 0.892 | 0.076 | -1.33 | 0.183 | 0.754 | 1.055 |
| rpt journal IF | 1.572 | 0.196 | 3.63 | 0.000 | 1.231 | 2.008 |
| rpt journal name | 1.053 | 0.141 | 0.38 | 0.703 | 0.809 | 1.369 |
| rpt pub total | 1.374 | 0.232 | 1.89 | 0.059 | 0.988 | 1.912 |

| Supplementary Table 14. Ordered logistic model predicting overall prestige of the journal/venue/publisher as a factor in publishing decisions (Model 8). Total n= 205. | | | | | | | | | | | | |
| --- | --- | --- | --- | --- | --- | --- | --- | --- | --- | --- | --- | --- |
| **Variable** | | **Odds Ratio** | | **Std Err** | | **z** | | **P value** | | **95% confidence interval** | | |
| age | | 0.895 | | 0.124 | | -0.80 | | 0.424 | | 0.682 | | 1.175 |
| gender | | 1.310 | | 0.369 | | 0.96 | | 0.338 | | 0.754 | | 2.275 |
| r-type | | 0.736 | | 0.220 | | -1.02 | | 0.305 | | 0.409 | | 1.323 |
| tenured | | 0.592 | | 0.209 | | -1.48 | | 0.138 | | 0.297 | | 1.183 |
| pubs published | | 1.093 | | 0.162 | | 0.60 | | 0.549 | | 0.817 | | 1.463 |
| rpt pub numbers | | 1.112 | | 0.190 | | 0.62 | | 0.534 | | 0.796 | | 1.554 |
| rpt preprint | | 1.180 | | 0.126 | | 1.55 | | 0.122 | | 0.957 | | 1.455 |
| rpt open access | | 0.827 | | 0.095 | | -1.64 | | 0.100 | | 0.660 | | 1.037 |
| rpt society | | 1.057 | | 0.091 | | 0.64 | | 0.524 | | 0.892 | | 1.252 |
| rpt journal IF | | 1.174 | | 0.142 | | 1.33 | | 0.182 | | 0.927 | | 1.487 |
| rpt journal name | | 1.420 | | 0.189 | | 2.64 | | 0.008 | | 1.095 | | 1.843 |
| rpt pub total | | 0.909 | | 0.154 | | -0.57 | | 0.572 | | 0.652 | | 1.266 |
| Supplementary Table 15. Ordered logistic model predicting public availability of the publication (i.e. open access) as a factor in publishing decisions (Model 9). Total n= 202. | | | | | | | | | | | | |
| **Variable** | **Odds Ratio** | | **Std Err** | | **z** | | **P value** | | **95% confidence interval** | | | |
| age | 1.103 | | 0.151 | | 0.72 | | 0.473 | | 0.843 | | 1.443 | |
| gender | 1.333 | | 0.367 | | 1.04 | | 0.296 | | 0.777 | | 2.287 | |
| r-type | 0.840 | | 0.258 | | -0.57 | | 0.569 | | 0.460 | | 1.532 | |
| tenured | 0.658 | | 0.222 | | -1.24 | | 0.215 | | 0.339 | | 1.275 | |
| pubs published | 1.115 | | 0.166 | | 0.73 | | 0.467 | | 0.832 | | 1.493 | |
| rpt pub numbers | 1.038 | | 0.170 | | 0.23 | | 0.820 | | 0.753 | | 1.431 | |
| rpt preprint | 1.033 | | 0.107 | | 0.31 | | 0.754 | | 0.843 | | 1.266 | |
| rpt open access | 1.957 | | 0.225 | | 5.84 | | 0.000 | | 1.562 | | 2.452 | |
| rpt society | 0.885 | | 0.082 | | -1.32 | | 0.185 | | 0.738 | | 1.061 | |
| rpt journal IF | 0.981 | | 0.114 | | -0.17 | | 0.869 | | 0.782 | | 1.231 | |
| rpt journal name | 1.129 | | 0.157 | | 0.87 | | 0.383 | | 0.860 | | 1.482 | |
| rpt pub total | 1.006 | | 0.164 | | 0.04 | | 0.971 | | 0.730 | | 1.386 | |

| Supplementary Table 16. Ordered logistic model predicting journal cost to publish as a factor in publishing decisions (Model 10). Total n= 192. | | | | | | |
| --- | --- | --- | --- | --- | --- | --- |
| **Variable** | **Odds Ratio** | **Std Err** | **z** | **P value** | **95% confidence interval** | |
| age | 0.866 | 0.124 | -1.01 | 0.315 | 0.653 | 1.147 |
| gender | 0.396 | 0.114 | -3.21 | 0.001 | 0.225 | 0.698 |
| r-type | 1.261 | 0.394 | 0.74 | 0.458 | 0.684 | 2.325 |
| tenured | 1.514 | 0.530 | 1.18 | 0.236 | 0.762 | 3.006 |
| pubs published | 1.159 | 0.174 | 0.99 | 0.324 | 0.864 | 1.555 |
| rpt pub numbers | 1.169 | 0.204 | 0.90 | 0.370 | 0.831 | 1.646 |
| rpt preprint | 1.113 | 0.119 | 1.00 | 0.317 | 0.903 | 1.372 |
| rpt open access | 1.041 | 0.114 | 0.37 | 0.713 | 0.840 | 1.290 |
| rpt society | 0.926 | 0.085 | -0.84 | 0.400 | 0.773 | 1.108 |
| rpt journal IF | 1.070 | 0.131 | 0.55 | 0.580 | 0.841 | 1.362 |
| rpt journal name | 0.850 | 0.115 | -1.20 | 0.231 | 0.652 | 1.109 |
| rpt pub total | 0.956 | 0.165 | -0.26 | 0.794 | 0.681 | 1.341 |
